# Coenzyme Q regulates UCP1 expression and thermogenesis through the integrated stress responses

**DOI:** 10.1101/2022.03.18.484957

**Authors:** Ching-Fang Chang, Amanda L. Gunawan, Peter-James H. Zushin, Greg A. Timblin, Ambre M. Bertholet, Biao Wang, Kaoru Sajio, Yuriy Kirichok, Andreas Stahl

## Abstract

Coenzyme Q (CoQ) is an essential component of mitochondrial respiration^1^ and required for thermogenic activity in brown adipose tissues^2^ (BAT). CoQ deficiency leads to a wide range of pathological manifestations^3^ but mechanistic consequences of CoQ deficiency in specific tissues such as BAT remain poorly understood. Here we show that pharmacological or genetic CoQ deficiency (50-75% reduction) in BAT leads to accumulation of cytosolic mitochondrial RNAs (mtRNAs) and activation of the eIF2α kinase PKR resulting in the induction of the integrated stress response (ISR) and suppression of UCP1 expression in an ATF4-dependent fashion. Surprisingly, despite diminished UCP1 levels, BAT CoQ deficiency increases whole-body metabolic rates at room temperature and thermoneutrality resulting in decreased weight gain on high fat diets (HFD). This mitohormesis like effect of BAT CoQ insufficiency is dependent on the ATF4-FGF21 axis in BAT revealing an unexpected role for CoQ in the modulation of whole-body energy expenditure with wide-ranging implications for primary and secondary CoQ deficiencies.

---

Coenzyme Q (CoQ) is a component of oxidative phosphorylation (OXPHOS) chain in mitochondria responsible for carrying electrons from multiple entry points to Complex III. While the functions of CoQ in the mitochondrial electron transport chain are well explored, basic questions about CoQ physiology such as its transport ^2^, participation in redox reactions outside of mitochondria^4^, and roles in mitochondrial to nuclear communications are still under investigated. CoQ deficiency has been associated with multiple human pathologies^1^. Primary CoQ deficiencies result from mutations in genes involved in CoQ_10_ biosynthesis. Secondary CoQ deficiency has been frequently detected in mitochondrial disorders, such as patients with mitochondrial DNA depletion syndromes^5^, and also can be linked to hydroxymethylglutaryl coenzyme A (HMG-CoA) reductase inhibitors, such as statins, which are commonly used in the treatment of hypercholesterolemia^6^. While the clinical benefits of CoQ supplementation remain controversial^7^, it is clear that statins have the ability to lower serum levels of CoQ^8^ and are associated with mitochondrial dysfunction in muscle^9^. In fact, CoQ deficiency commonly affects organs rich in mitochondria and with high energy demand^1^ such as heart, brain and kidney. Brown adipose tissue (BAT) facilitates nonshivering thermogenesis through the actions of uncoupling protein 1 (UCP1), which allows for the dissipation of metabolic energy in the form of heat, and features highly active and densely packed mitochondria^10,11^. BAT has drawn much attention due to its high metabolic rates upon activation making it a promising target for anti-obesity and antidiabetic interventions^12^. We previously showed that BAT CoQ uptake is dependent on the scavenger receptor CD36 and demonstrated that the loss of CD36 results in cold-intolerance^2^. However, the molecular mechanisms by which CoQ deficiency triggers thermogenic dysfunction is still not fully understood.

Mitochondrial stress, such as dysfunction in OXPHOS, can trigger an evolutionarily conserved transcriptional program called the mitochondrial unfolded protein response (UPR^mt^) which in turn leads to the activation of the integrated stress response (ISR). ISR suppresses general translation via the phosphorylation and inactivation of the eukaryotic translation initiation factor 2 (eIF2) complex subunit 2α (eIF2α) by upstream stress induced eIF2α kinases^13-15^. However, ISR allows for the selective production of proteins such as ATF4 from a subset of mRNAs which frequently possess multiple upstream open reading frames^16^. UPR^mt^, mostly studied in *C. elegans*, can be triggered by improper mitochondrial protein import capacity resulting in compensatory changes to restore mitochondria homeostasis. Interestingly, activation of adaptive responses to mitochondrial stress can provide metabolic benefits^17^ and promote longevity^18^ in a process termed mitohormesis^19^. In mammals, components of UPR^mt^ signaling are still under investigation and appear to be tightly integrated with the ISR^14^. Thus, UPR^mt^ activation in response to mitochondria stressors in mammals results in the selective translation of ISR-related transcription factors, such as activating transcription factors ATF4, ATF5 and DNA-damage-inducible transcript 3 (CHOP). How CoQ insufficiency affects UPR^mt^/ISR in mammalian cells remains to be explored.

## CoQ deficiency changes BAT mitochondrial respiration via suppresses UCP1

In order to investigate the impact of CoQ deficiency on thermogenic function in brown adipocytes, we used the polyprenyl-diphosphate:4-HB transferase (COQ2) inhibitor 4CBA to block CoQ biosynthesis^20^. Total CoQ levels, dominated by CoQ_9_ in murine cells, were reduced to 40-50% by 4CBA in mature brown adipocytes from an immortalized preadipocyte cell line^21^ (Fig. 1A). CoQ is known as a critical component for mitochondrial respiration, thus we examined cellular respiration of CoQ deficient brown adipocytes using a standard mitochondrial stress test (Fig. 1B). Interestingly, while basal and uncoupled respiration were affected by 4CBA treatment, ATP-linked respiration was unchanged (Fig. 1C). Expression of mitochondrial electron transport chain complexes was also unaffected by 4CBA treatment (Fig. S1A). Surprisingly, CoQ deficiency triggered a robust suppression of *UCP1* gene and protein expression and downregulated several key brown/beige transcriptional regulators including *PGC1*α, *EBF2* and *PPAR*γ, but not *PRDM16* (Fig. 1D-E). This suggests that the loss of UCP1, rather than defective electron transfer, dominates the observed OXPHOS decrease. Exogenous CoQ_10_ supplementation of CoQ deprived cells significantly improved basal and proton leak respiration, and UCP1 expression (Fig. 1B and S1B). Patch-clamp analysis (Fig. S1C) showed that the uncoupling function of UCP1 was independent of CoQ supporting the notion that primary defects are due to changes in UCP1 abundance but not function. To examine BAT CoQ deficiency in vivo, we generated a brown/beige fat-specific knockout of the CoQ synthesis enzyme decaprenyl diphosphate synthase subunit 2 (PDSS2) by crossing UCP1-cre animals with a PDSS2 Flox strains, both in the C57 background, generating the PDSS2^BKO^ line. PDSS2^BKO^ animals display abrogation of *PDSS2* expression in BAT and a significant (75%) reduction of CoQ levels in that, but not other, tissues while plasma CoQ levels were increased (Fig. 1F-G). Importantly, a pronounced suppression of *UCP1* was detected in BAT tissues of the PDSS2^BKO^ animals (Fig. 1G). UCP1 suppression was also observed more acutely *in vitro* using *PDSS2* knockdown brown adipocytes (Fig. 1H), demonstrating that both genetic and pharmacological BAT CoQ deficiency result in robust UCP1 suppression.

**Fig. 1.**
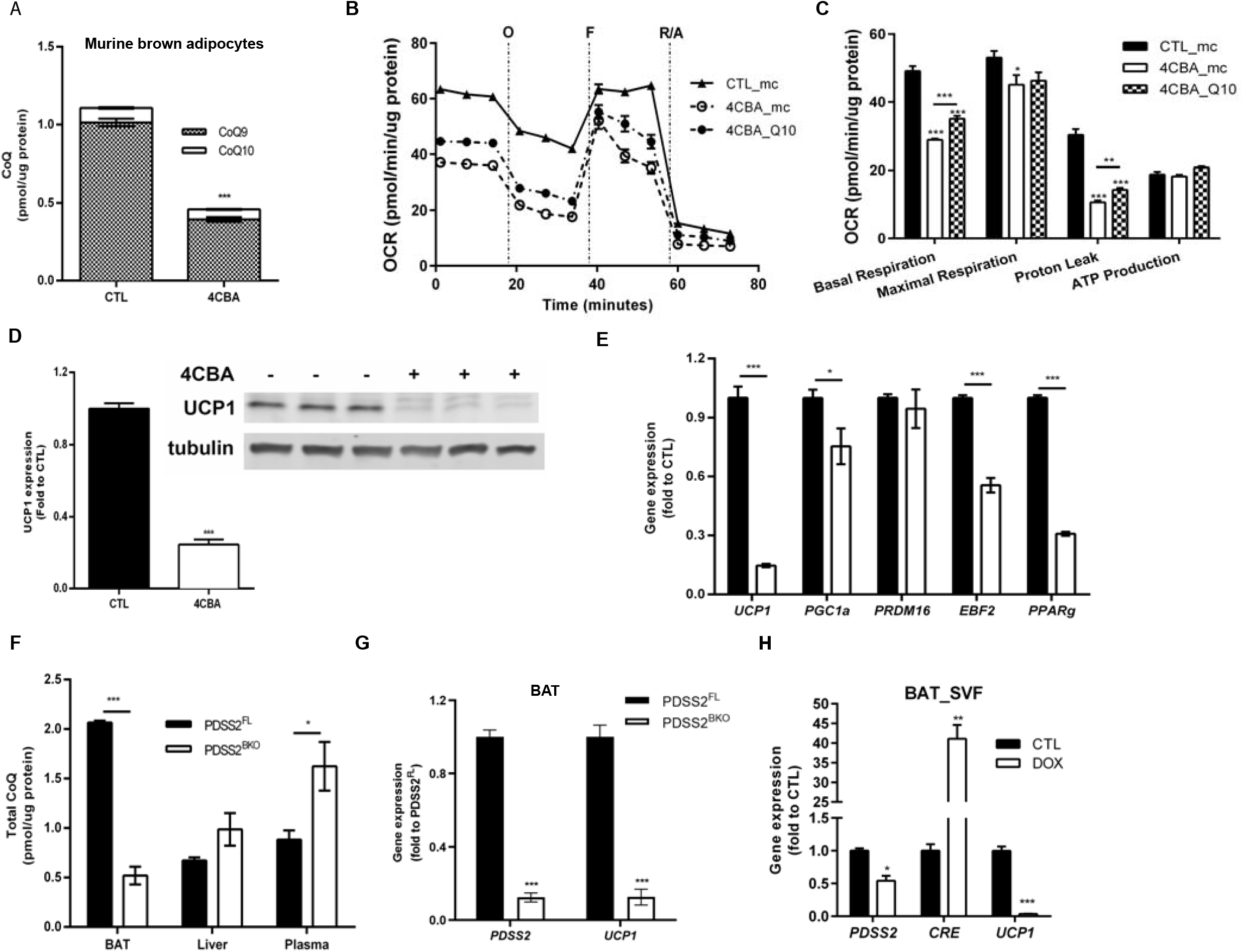
UCP1 downregulation by CoQ deficiency (A) Coenzyme Q (CoQ) levels in murine brown adipocytes treated with vehicle (CTL) or 4CBA, n=3. (B-C) Oxygen consumption rate (OCR) was measured using seahorse XFe24 analyzer and components of mitochondrial respiration were analyzed, n=5. (D) UCP1 protein expression, n=3 (E) Thermogenic genes were measured by rtPCR, n=6. (F) CoQ levels in tissues of BAT-specific PDSS2 knockout mice (PDSS2^BKO^) and controls (PDSS2^FL^). (G) gene expression in BAT measured by rtPCR, n=7-10. (H) Stromal vascular fraction (SVF) was isolated from BAT of PDSS2^FL^ animals and PDSS2 was knockout by the induction of cre recombinase with doxycycline. Gene expression was measured by rtPCR (n=3). Data are mean ± SEM, representative from three repeated experiments.

## BAT CoQ deficiency activates ISR and UPR^mt^

To further define the transcriptional signature of CoQ deficiency, we performed RNA sequencing (RNAseq) analysis of mature brown adipocytes treated with 4CBA or vehicle for 48 hours. Gene Ontology (GO) analysis identified enrichment in mitochondrial dysfunction for categories including TCA cycles, respiratory electron transport and lipid catabolic process repressed by 4CBA (Fig S2A). Key components identified in endoplasmic reticulum and mitochondrial unfolded protein responses (UPR) were increased by 4CBA (Fig .2A and S2B). CoQ deficiency-mediated mitochondria dysfunction was also validated with increases of elongated mitochondria and decreased phosphorylation of dynamin-related protein 1 (Drp1) at serine 616, the active form which facilitates mitochondrial fission (Fig. S2C-D), further indicating reduced thermogenic activities^22^. Differential expression analysis confirmed *UCP1* to be among the genes significantly reduced in expression. Up-regulated genes included notable ISR components such as *ATF4, ATF5, CHOP*, and *FGF21* (Fig. 2B). HOMER motif analysis also revealed CHOP, ATF4 and C/EBP binding motif, key UPR^mt^ and ISR effectors^23,24^, as top enriched transcription factor binding motifs in the promoters of the significantly upregulated genes (Fig. 2C). To determine the temporal aspects of ISR induction by CoQ deficiency, we performed rtPCR assays after 7, 24, 48, and 72h of 4CBA treatment. Induction of *ATF3, -4*, and *-5* was rapid, peaking between 7 to 24h, and stayed significantly elevated above baseline (Fig. S2E). Similarly, UPR^mt^ associated genes such as *HSP60, LONP1*, and *CLPP* rose rapidly, peaking at 24h and subsequently declined (Fig. S2F). In contrast, ISR related gene^13^ *FGF21, GDF15, MTHFD2*, and *PHGDH* also rose rapidly at 7h but stayed robustly elevated throughout all treatment time points (Fig. S2G) indicating a well-orchestrated cellular response to CoQ depletion that results in compensatory changes. ISR is an adaptive translation program activated by different stress signals that leads to the phosphorylation at the α subunit of the initiation factor eIF2 and inactivation of cap-dependent translation^23^. CoQ deficiency induced eIF2 phosphorylation concomitant with decreased UCP1 expression, and induction of UPR^mt^ and ISR mediators was observed in 4CBA treated brown adipocytes (Fig. S2E-G and Fig. 2D) as well as in vivo in the BAT of PDSS2^BKO^ animals (Fig. 2E-F).

**Fig. 2.**
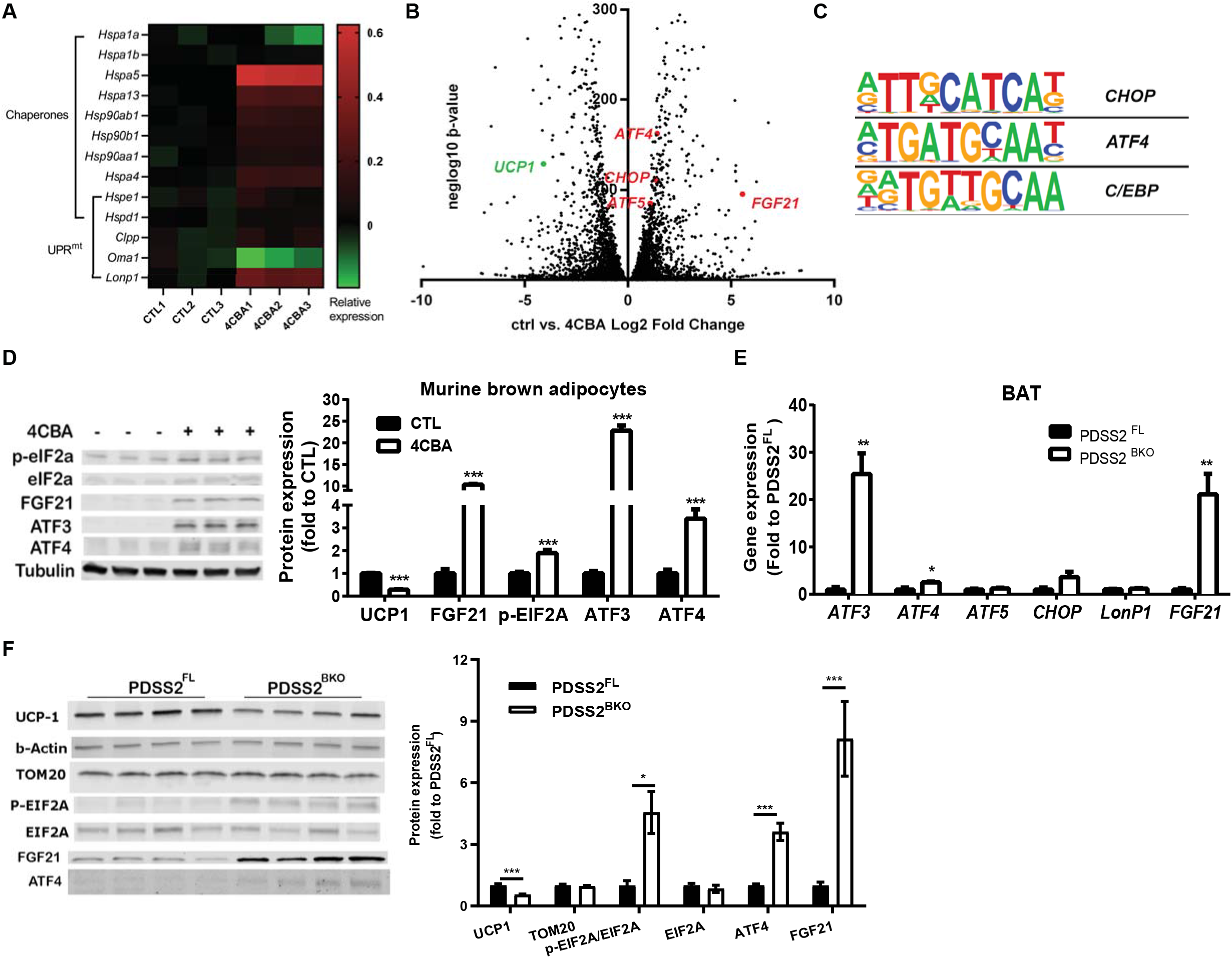
ISR and UPR^mt^ are triggered by CoQ deficiency (A) Relative expression (log_10_ -transformed RPKM value) of UPR-associated genes in murine brown adipocytes treated with 4CBA (48h). (B) Volcano plot obtained from DESeq2 analysis of vehicle control and 4CBA-treated murine brown adipocytes RNA pools, n=3. (C) The top 3 enriched transcription factors identified by Motif analysis, P<1e-6. (D) ISR-related protein levels in brown adipocytes after 48h treatment. (E-F) Genes or proteins expression in BAT of PDSS2^FL^ or PDSS2^BKO^ animals, n=4. Data are mean ± SEM.

## BAT CoQ deficiency triggers cytosolic accumulation of mitochondrial RNA, PKR activation, and ATF4 induction

To identify the ISR effectors responsible for UCP1 suppression as the consequence of CoQ deficiency, we attempted to rescue UCP1 expression in CoQ-deficient brown adipocytes via knockdown of ATF4, ATF5, and CHOP. siRNA mediated knockdown efficiency was comparable at 50% (Fig. S3A). While knockdown of ATF5 or CHOP alone only increase 15-25% *UCP1* expression in 4CBA treated cells (Fig. S3B), ATF4 knockdown rescued ∼30% of UCP1 expression, which could be further increased by simultaneous CHOP, but not ATF5, knockdown (Fig. 3A), suggesting an involvement of an ATF4/CHOP heterodimer in UCP1 regulation in response to ISR.

**Fig. 3.**
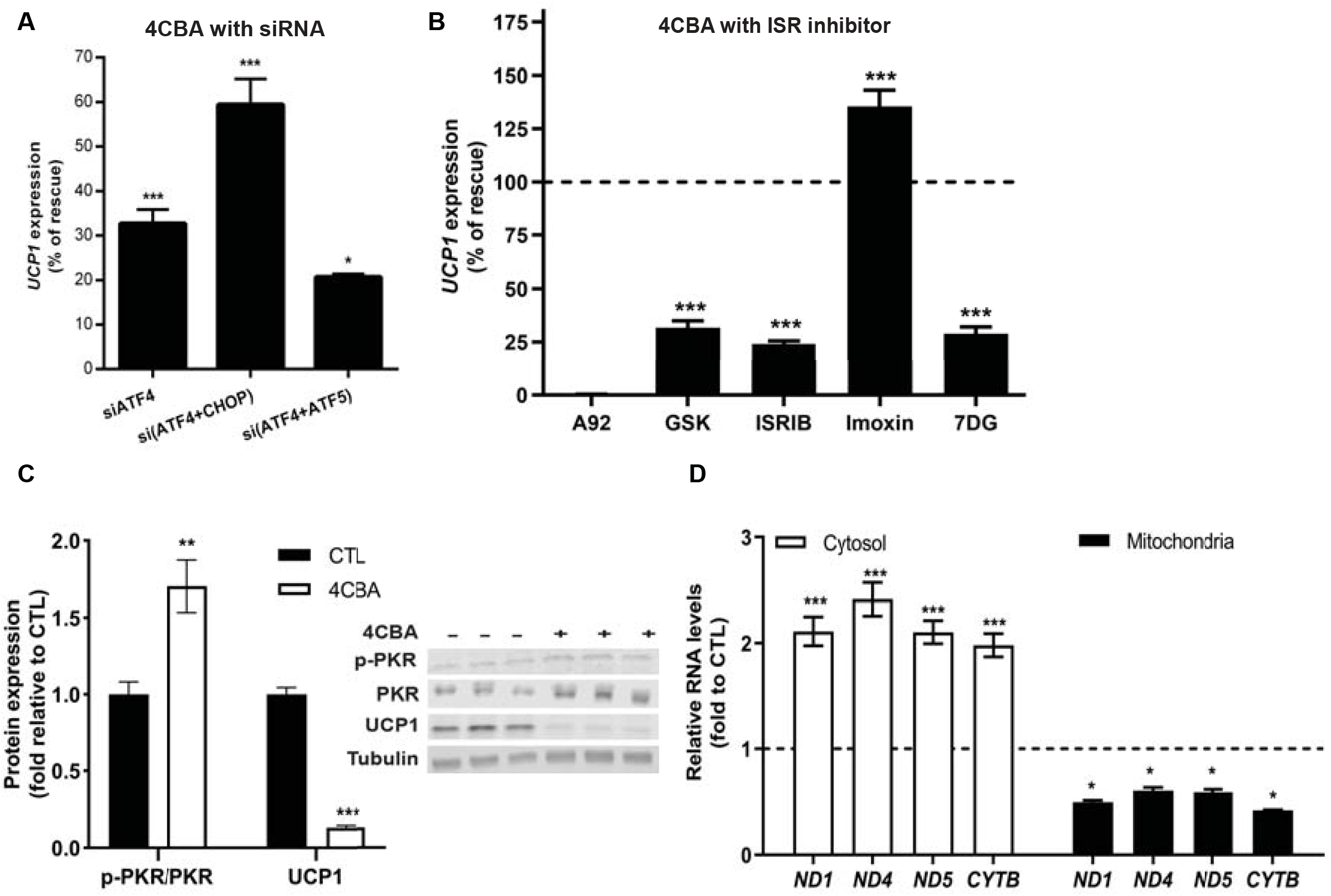
UCP1 suppression by CoQ deficiency depends on ISR (A) *UCP1* gene expression in brown adipocytes with siRNA knockdown of ATF4, ATF4 and CHOP or ATF4 and ATF5 following 24h-4CBA treatment. Data represents % of rescue. *UCP1* inhibition by 4CBA at scramble siRNA control was set as 0% rescue. (B) UCP1 expression in brown adipocytes treated with 4CBA in the presence of ISR inhibitors. (C) PKR and phosphorylated PKR (Thr451) levels. (D) Mitochondrial transcript levels in cytosol or mitochondria; Data are mean ± SEM; n=4-6/group.

The core event in the ISR is the phosphorylation of the α subunit of eIF2 at ser51 by members of eIF2α kinase: PERK, GCN2, HRI and PKR^25^. To better understand the upstream effects triggering UCP1 suppression in CoQ-mediated ISR, we treated cells with 4CBA and pharmacological inhibitors of eIF2α kinases or other components of the ISR^23^ pathway including PKR inhibitors imoxin and 7DG, GCN2 inhibitor A92, PERK inhibitor GSK2606414 (GSK) and ISRIB. While the GCN2 inhibitor A92 failed to rescue the *UCP1* suppression phenotype, the PERK inhibitors GSK, the reversible PKR inhibitor 7DG and ISRIB had mild, but significant effects, in 4CBA treated cells (Fig. 3B). It has been shown HRI is required for ISR activation upon mitochondria stress^26^. We used siRNA targeting HRI to examine the involvement of HRI in ISR dependent UCP1 suppression by CoQ deficiency (pharmacological inhibitors are unavailable). While siRNA treatment resulted in a robust 60% reduction of HRI transcripts, it did not rescue UCP1 expression significantly (Fig. S3C). Interestingly, treatment of CoQ deficient cells with the PKR inhibitor imoxin robustly rescued *UCP1* expression (Fig. 3B) indicating a key role for this eIF2α kinase in the ISR activation by CoQ deficiency. Activation of PKR subsequent to 4CBA treatment in murine brown adipocytes was further supported by increased Thr451 phosphorylation (Fig. 3C). PKR can be activated by viral RNAs but also by cytoplasmic mitochondrial double-stranded RNA (mtRNAs) ^27^. We found that levels of mitochondrial transcripts of ND1, ND4, ND5 and CYTB were increased in the cytoplasmic fraction, but decreased in the mitochondria of the 4CBA-treated brown adipocytes (Fig. 3D). This suggest that mitochondrial CoQ deficiency induces the ISR via increased mtRNAs export and PKR-mediated eIF2α phosphorylation.

## BAT CoQ deficiency modulates whole body metabolism

To further examine how BAT CoQ deficiency impact whole body metabolism, we explored the metabolic alterations on BAT-specific PDSS2 knockouts using CLAMS metabolic chambers at thermoneutrality (30°C) and mild cold challenges (23°C). We were surprised to observe that core body temperature was normal (data not shown) and that VO_2_ and VCO_2_ rates during the light cycle (low activity) and dark cycle (high activity) were increased in PDSS2^BKO^ animals compared to PDSS2^FL^ controls. This held true both at room temperature, a mild cold challenge for mice, and at thermoneutrality when BAT respiration should be suppressed (Fig. 4A-B). Lower ambulatory activity was measured in the PDSS2^BKO^ animals compared to PDSS2^FL^ controls (Fig. 2C) indicates the increased respiration is irrelevant to physical activity. However, when shifting the animals to 4°C, an immediate and dramatic respiration defect became apparent with significantly lower VO_2_ rates after 2h of cold exposure and became hypothermic between 4-5 hours at 4°C (Fig. S4A). While no significant difference was observed in food intake among the genotypes (Fig. S4B), the respiratory exchange ratio (RER) was higher (0.95 compared to 0.88 for PDSS2^FL^) in PDSS2^BKO^ at 23°C (data not shown). In line with enhanced metabolic rates, PDSS2^BKO^ animals also gained significantly less body weight than control littermates particularly when animals were fed with 60% fat diets for 12 weeks (Fig. 4D-E), suggesting the metabolic alterations by CoQ deficiency in BAT provide protection from diet-induced weight gains.

**Fig. 4.**
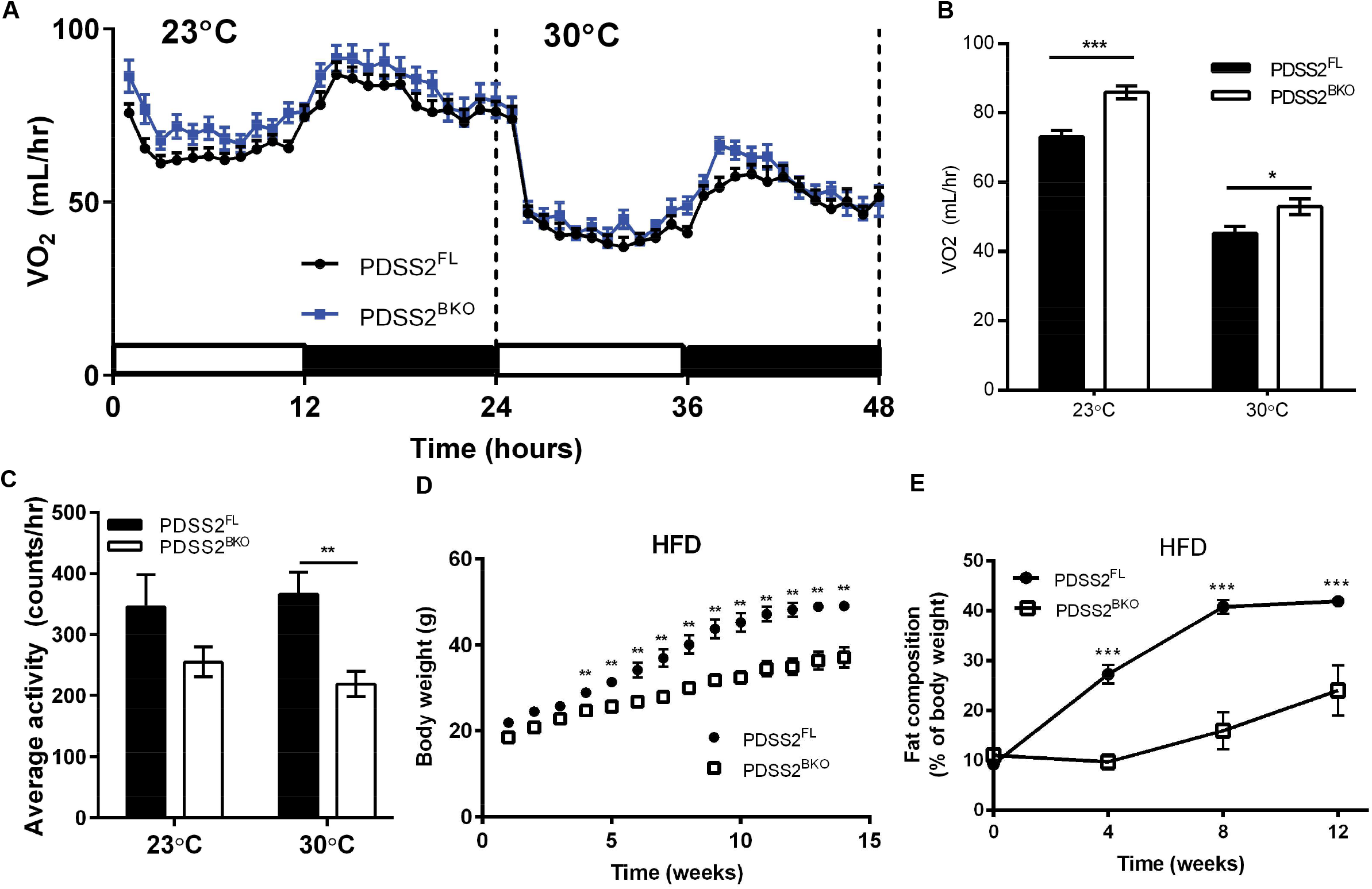
CoQ deficiency leads to increased metabolic rates in *in vivo* model (A) Oxygen consumption rate of PDSS2^FL^ or PDSS2^BKO^ animals was monitored under 23 and 30°C in real-time in metabolic cages using CLAMS. (B) averaged oxygen consumption rate. (C) Average ambulatory activity in 24 hours at 23 and 30°C. (D, E) Body weight and fat compositions were monitored during 12 weeks of HFD feeding; Data represents mean ± SEM, n=10-11/group (A-C) or 4/group for HFD.

## ATF4-FGF21 axis is crucial in metabolic adaptation by BAT CoQ-deficiency

In addition to its thermogenic function, BAT is also an endocrine tissue secreting several important metabolism regulating hormones including FGF21^28^. Enhanced expression of FGF21 has been linked to activation of the ISR pathway^13^, and can induce UCP1-independent thermogenic processes in adipose and other tissues^29,30^. Interestingly, not only did 4CBA treatment rapidly induced a 30 -to -40-fold increase in FGF21 transcripts in brown adipocytes in tissue culture (Fig. S2G), increase in FGF21 was also observed in BAT of normal- or HFD-fed PDSS2^BKO^ animals (Fig. 2E and S5A). Enhanced FGF21 expression by BAT also translated to 4-5-fold increase in circulating FGF21 levels (Fig. 5A-B). FGF21 expression remained unchanged in other tissues (Fig. S5B) including the liver, which is considered the main site of FGF21 production^31^, or inguinal white adipose tissue (iWAT), suggesting the observed elevation of FGF21 in circulation was mainly secreted by the BAT (Fig. S5C), and the enhanced FGF21 secretion is part of the compensatory mechanisms allowing CoQ deficient animals to cope with decreased UCP1 expression. Moreover, the ISR markers were not elevated in the livers of liver-specific PDSS2 knockout animals (PDSS2^AlbKO^) (Fig. S5D), indicating the ATF4-FGF21 axis in ISR induced by CoQ deficiency is tissue-specific.

**Fig. 5.**
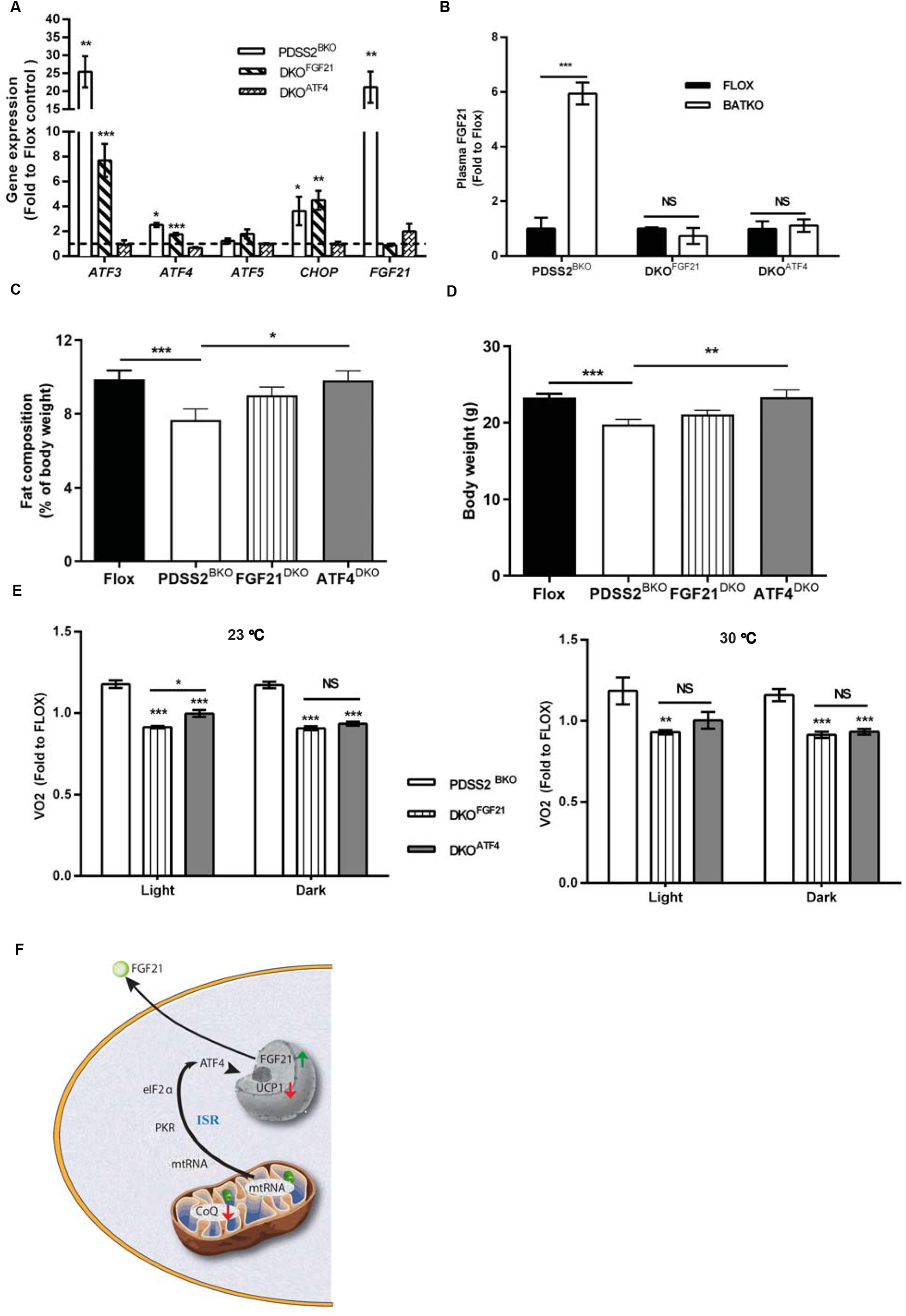
Metabolic impacts by ISR in CoQ-deficient BAT (A) ISR-related gene expression in BAT of PDSS2^BKO^ or BAT-specific PDSS2 and FGF21 or ATF4 double knockouts (B) Plasma FGF21 levels of BAT specific PDSS2KO (PDSS2^BKO^), double knockouts DKO^FGF21^ or DKO^ATF4^. (C-D) Body weight and fat content of control Flox, PDSS2^BKO^ and double knockouts (DKO^FGF21^, DKO^ATF4^) after 2-week defined diet at age 10-week. VO_2_ of PDSS2^BKO^, DKO^FGF21^ or DKO^ATF4^ at 23°C or 30°C. Data represents mean ± SEM, n=6-7/group. (F) Graphic scheme illustrated the mechanisms of mitohormetic responses by CoQ deficiency in BAT.

To evaluate how ATF4-FGF21 axis in CoQ-deficient BAT influences whole-body metabolism in PDSS2^BKO^ animals, we generated BAT-specific PDSS2 and FGF21 or PDSS2 and ATF4 double knockouts (DKO^FGF21^ or DKO^ATF4^) by crossing our PDSS2^BKO^ and FGF21 or ATF4 Flox animals. BAT from DKO^FGF21^ showed an increase in ISR related genes *ATF3, -4, -5* and *CHOP*, like those observed in BAT of PDSS2^BKO^. The induction of *FGF21* expression in BAT of DKO^FGF21^ animals was blunted (Fig. 5A) and was reflected in the circulating level of FGF21 (Fig. 5B). Decreased adiposity of the PDSS2^BKO^ animals was normalized in the DKO^FGF21^ animals, as there was no significant difference in body or adipose tissue weight as measured by EchoMRI (Fig. 5C-D), without any influence on food intake (data not shown). VO_2_ rates during the light cycle and dark cycle were significantly decreased in DKO^FGF21^ animals compared to PDSS2^FL^ controls or PDSS2^BKO^ animals in either room temperature (23°C) or in thermoneutrality (30°C) (Fig. 5E-F). Additionally, BAT-specific dual ATF4/PDSS2 knockout animals DKO^ATF4^ also abolished the increased plasma FGF21 (Fig. 5B) and VO2 (Fig. 5E-F) confirming the regulatory role of the ATF4-FGF21 axis on metabolic regulation by BAT CoQ deficiency. Interestingly, we also found upregulation of genes in mechanisms of UCP1-independent thermogenesis in BAT of PDSS2^BKO^ animals, including *SLC6A8* and *GATM* in creatine-dependent thermogenesis^32^, *Ryr2* and *SERCA2* in calcium cycling^33,34^, and mitochondrial ADP/ATP carrier *AAC2* in AAC-mediated H^+^ leak^35^ (Fig. S5E). The activation of UCP1-independent thermogenesis remained in BAT-specific dual FGF21/PDSS2 (DKO^FGF21^), but it was abolished at ATF4/PDSS2 (DKO^ATF4^) knockout animals (Fig S5E), emphasizing the regulatory roles of ATF4 on the alternative thermogenesis in CoQ deficient BAT. On the other hand, induction of *AAC2* was also found in iWAT of PDSS2^BKO^ animals, but was abolished in double knockouts (Fig. S5F), suggesting FGF21’s regulatory role on alternative thermogenesis in iWAT of PDSS2^BKO^ animals and potentially through endocrine route from BAT. Our data indicates circulating FGF21 secreted from CoQ-deficient BAT provided systemic metabolic effects which maintains PDSS2^BKO^ animals a lean phenotype, with ATF4-mediated ISR signaling plays a central role for the observed metabolic adaptation.

## Discussion

CoQ deficiency can result from genetic mutations in the biosynthesis pathway (primary CoQ deficiency), from mitochondria disorders, or pharmacological inhibition of CoQ synthesis (secondary CoQ deficiency). Secondary CoQ deficiency is more common as it is a side-effect of the widely used cholesterol lowering HMGCoA reductase inhibitors (frequently referred to as statins). The effects of CoQ deficiency on specific tissues are still ill understood but are thought to disproportionately affect cells with high energy demands. Based on the classical roles of CoQ in the ETC, BAT CoQ deficiency would have been predicted to result in decreased maximal mitochondrial and whole-body respiration and enhanced susceptibility to HFD challenges due to decreased caloric output. Instead, we found that BAT CoQ deficiency enhances whole-body respiration, which we suggest is downstream of a mechanistic cascade that links CoQ insufficiency in BAT with the appearance of cytosolic mtRNA, PKR activation, induction of ISR/ATF4, suppression of UCP1, but also massive induction of FGF21 expression and secretion which ultimately results in mitohormesis like response (Fig 5F).

Human CoQ deficiencies caused by either genetic or pharmacological inhibition of CoQ synthesis have highly variable clinical presentation and differential impacts on multiple organs, and the underlying molecular mechanisms are complicated and still unknown. Clinical studies showed human CoQ deficiency displays CoQ_10_ reduction of 40-85 % in skin fibroblasts, muscles, blood mononuclear or plasma^36,37^; However, thus far, muscle biopsy is the most common clinical method to detect primary CoQ deficiency and CoQ levels in adipose tissues of primary CoQ deficiency patients are not available. We used pharmacological and genetic BAT CoQ deficiency mouse models here to examine how CoQ deficiency alters BAT function and potential effects on whole body metabolism. Complete loss of CoQ in tissues with primary CoQ synthesis defects is apparently avoided as exogenous dietary CoQ can partially compensate^38^. We found 50% residual CoQ in the PDSS2 knockout BAT which can be further reduced to 25% by limiting dietary CoQ intake. Interestingly, CoQ deficiency and mitochondrial dysfunction in *C. elegans* led to reduction in mitochondrial respiration, but stimulated cell-protective responses and increased life span through activation of the UPR^mt^ pathway in the key tissues (intestines and neurons) in a cell non-autonomous manner^39^. In mouse models, the physiological manifestations of CoQ deficiency depend on tissue and cell types. Kidney and cerebellum are the most often affected organs in primary CoQ deficiency while liver appears to be insensitive to it^3^. Interestingly, BAT CoQ deficiency triggered UPR^mt^ and ISR which resulted in increased basal metabolic rates and amelioration of diet-induced obesity, independent of interrupting mitochondrial respiration. These effects were not observed in a simultaneously generated liver-specific CoQ deficiency animal strain (PDSS2^AlbKO^), indicating the UPR^mt^ /ISR induction by CoQ deficiency presents a tissue-specific manner.

In mammalian cells disruption of mitochondrial functions is closely coupled to activation of ISR, which can be triggered by four different eIF2α kinases, PKR, GCN2, HRI and PERK^14,26,40^. It has been shown that mitochondrial RNAs can activate PKR under stress conditions^41,42^ and activation of PKR was associated with inflammation and metabolic disorders^43^. In line with these findings, we observed accumulation of cytosolic mtRNAs exported from mitochondria acted as the upstream ISR activator on PKR, and the ISR effector ATF4 play critical roles in UCP1 suppression by CoQ-deficiency in BAT. Although other eIF2α kinase GCN2, HRI, or PERK have been linked to triggering ISR activation in response to mitochondrial dysfunction^26,44^ or key regulator in ER-mitochondria communication and mitochondria homeostasis ^45^, we were unable to reverse UCP1 suppression by CoQ deficiency in brown adipocytes via genetic or pharmacological inhibition of these kinases. Together, this suggests that PKR plays a major role in ISR activation in CoQ deficient brown adipocytes. Increased cytosolic mtRNAs as a result of mitochondrial CoQ deficiency could act as the PKR activator to regulate ISR pathways.

Regarding the regulation of UCP1 expression by the ISR, it is noteworthy that UCP1 and PGC1a expression is increased in the BAT of ATF4 whole body knockout mice^46^, suggesting that ATF4 can act as a negative regulator of uncoupled respiration. Transcription of UCP1 can be positively or negatively controlled by several transcription factors, such as members of the cAMP response element-binding protein (CREB) family, the peroxisome proliferators-activated receptors (PPAR), the CCAAT/enhancer-binding proteins (C/EBP), and liver X receptor^47,48^. However, it remains to be seen if ATF4 can directly or indirectly bind on regulatory elements in the UCP1 gene and repress UCP1 transcription. Importantly, while a previous publication had suggested a role of CoQ as an obligatory cofactor for UCP1^49^, we were unable to show an effect of CoQ on proton conductance by UCP1 using a mitoplast patch-clamp technique indicting that the reduction of uncoupled respiration resulting from CoQ deficiency is due to reduced UCP1 expression and not impaired function.

The stress hormone FGF21 is the master metabolic regulator in upregulation of energy expenditure and has promising therapeutic application for the treatment of obesity-related metabolic disorders^50^. Liver is thought to be the main organ for FGF21 production, but it can be synthesized and secreted from other tissues during stress responses as well ^13^. FGF21 is transcriptionally regulated by ATF4 through three binding sites of Amino Acid Response Element (AARE) in its promoter region^51^. We found that thermogenic fat BAT, rather than liver, - derived FGF21 was responsible for the increased energy expenditure in both mild cold (23°C) and thermoneutrality (30°C) conditions, which contributed to a leaner body composition. Additionally, deletion of upstream ATF4 or the effector FGF21 in CoQ-deficient BAT abolished the compensatory UCP1-independent thermogenesis and metabolic modifications confirming the physiological role of an ATF4-FGF21 axis on CoQ deficient disease model in BAT. In agreement with our study illustrating the essential roles of ISR in metabolic adaptation, recent reports also showed the ATF4 is required for metabolic adaptation in the BAT-specific OPA1 and BAT-specific Lrpprc knockout animals^52,53^, We cannot exclude other metabolic modulators, such as GDF15 or small extracellular vesicles^54^, secreted from CoQ-deficient BAT activate inter-organ mitohormesis and contribute to the systemic metabolic alteration.

Overall, the unexpected consequences of BAT CoQ deficiency, including a seemingly paradoxical anti-obesity response, are likely to have important implications for our understanding of CoQ related disorders and as yet undescribed BAT-dependent effects of statin therapy.

## Methods

### Mouse model

All animal experiments were performed under the guidelines established by the UC Berkeley Animal Care and Use Committee. Brown fat specific PDSS2 knockout PDSS2^BKO^ were generated by crossing PDSS2-LoxP mice (gift from Dr. Gasser^3^) with UCP1-cre mice (mouse model developed by the Rosen lab deposited in JAX, stock #024670) on a C57BL/6J background. Liver specific PDSS2 knockouts (PDSS2^AlbKO^) were also generated by crossing PDSS2-LoxP mice with Albumin-cre mice (^55^; JAX, stock # 003574). Animals were fed regular diet (LabDiet 5053) under the ambient temperature at 23°C. Male animals aged between 6-8-week-old were fed defined diet (Research diet D12450J) for two weeks before experiments, unless described else. The defined diet contains only 32% of CoQ content compared to normal chow thus is suitable in our experiments for limiting CoQ intake. Diet induced obesity was performed on male animals at age of 6 week fed with 60 kcal% fat diet (Research diet D12492) for 12 weeks. Food consumption was monitored once per week and accumulated food consumption was shown as accumulated amount over two weeks per animals. PDSS2 flox allele control animals (PDSS2^FL^) were used as control relative to PDSS2^BKO^. Brown fat specific PDSS2 and FGF21 double knockout DKO^FGF21^ were generated by crossing PDSS2^BKO^ with FGF21-LoxP mice^56^ (JAX stock #022361). Brown fat specific PDSS2 and ATF4 double knockout DKO^ATF4^ were generated by crossing PDSS2^BKO^ with ATF4-LoxP mice^57^. Flox/Flox littermates were used as control, respectively. ALL animal studies were performed using age-matched male mice (6-12 weeks), cohorts of greater than or equal to six mice per genotype or treatment, and repeated at least twice.

### Lentiviruses

Doxycycline inducible lentiviral Cre recombinase expression plasmid pLenti-Cre was constructed by replacing the complete attR1-CmR-ccdB-attR2 of the destination vector pInducer20, a gift from Stephen Elledge ^58^ (Addgene plasmid # 44012 ; RRID:Addgene_44012), with the cre recombinase gene of the entry vector pENTR1a-cre (a gift from Xin Chen at UCSF) using Gateway cloning technique (Invitrogen).

Lentiviruses were produced using Lipofectamine3000 (Invitrogen) according to the manufacturer’s instructions. Briefly, 293T cells were transfected with pLenti-cre, with psPAX2 (a gift from Didier Trono, Addgene plasmid # 12260; RRID:Addgene_12260), pMD2.G (gift from Didier Trono, Addgene plasmid # 12259 ; RRID:Addgene_12259) in Lipofectamine3000. Supernatants were harvested at 24h and 48h later, clarified, filtered through a 0.45-μm filter, and stored at -80°C.

### Cell cultures

Murine brown/beige cells: Immortalized murine brown preadipocyte line is a generous gift from Kajimura Lab^21^. The SVFs were isolated from BAT fat depots of either C57/B6 or PDSS2^FL^ male mice aged 8-12 weeks. Cells were cultured in DMEM supplemented with 10% FBS, penicillin, and streptomycin. Murine brown adipocytes were prepared from adipogenic differentiation of the confluent preadipocytes/SVFs with DMEM containing 10% FBS, penicillin/streptomycin and differentiation cocktails (850 nM insulin, 2 nM T3, 125 nM indomethacin, 0.5 mM 3-isobutyl-1-methylxanthine, 1μM dexamethasone, and 1 μM rosiglitazone) for 2 days. Two days after induction, cells were switched to a maintenance medium containing 10%FBS, P/S, insulin, T3 and rosiglitazone. Experiments assessing CoQ deficiency on thermogenic function were performed on day 4-6 of differentiation (48h) or day 5-6 (24h) in the presence of 4-chlorobenzoic acid (4CBA, 4mM) or vehicle (0.1% DMSO and 1% ethanol in culture medium) for murine adipocytes. Primary beige adipocytes were treated at day 6-8 based when the expression of adipocyte markers FABP4 and UCP1 reached plateau (indicators of adipocyte maturation), n=3 independent biological replicates per conditions. ISR inhibitors were added with 4CBA (4 mM) in the maintenance medium at day 5 of differentiated adipocytes for 24 hours for assessment of UCP1 rescue effects. ISRIB (200 nM), GSK (1 μM), A92 (10 μM), Imoxin (2 μM), or 7DG (10 μM).

Lentivirus infection: Isolated SVFs from BAT of PDSS2^FL^ male mice were cultured in DMEM supplemented with 10% FBS, penicillin, and streptomycin. Cells were then incubated with medium containing lentivirus (pLenti-cre construct) for 16-18 hours. Adipogenic differentiation was initiated once cells reached full confluency as previously described. After 5 days of adipogenic differentiation, the adipocytes derived from SVFs of PDSS2^FL^ male mice were switched to a DMEM medium supplemented with doxycycline (0.5 ug/ml) and 10% lipoprotein free serum for 48 hours.

### CoQ_10_ supplementation

CoQ_10_ loaded micelles were prepared as previously published^59^. Briefly, Kolliphor HS15 (30mg) and CoQ_10_ (5 mg) were dissolved at 50 °C with constant stirring, and 3 ml of glucose solution (0.2% w/v) containing sodium chloride (0.1538 M) was added in drop by drop while stirring. CoQ_10_ concentration in the micelle solution was determined by HPLC method. Final concentration 2 μM of the micelle formulation CoQ_10_ was used in the described experiments.

### Coenzyme Q extraction and measurement

CoQ levels from tissues cells, or isolated mitochondria were extracted using hexane as previously described^2^. Quantification of CoQ_9_ and CoQ_10_ was determined using Agilent 1100 HPLC system on a C18-ODS Hypersil reverse phase column (Thermo Scientific #30105-254630). Samples were spiked with CoQ_4_ to calculate the extraction and recovery efficiency. The CoQ analysis was performed with a flow rate of 1 mL/min with a mobile phase consisting of solution A (80% methanol/20% water) and solution B (100% ethanol) for 40 min. CoQ4, CoQ9 and CoQ10 peaks appear around 8 min, 23.8 min and 24.6 min, respectively. Absolute CoQ amounts was determined using a standard curve generated from commercial CoQ_9_ and CoQ_10_ (Sigma).

### Mitochondria isolation

Mitochondria was isolated as previously described^2^. Briefly, murine brown adipocytes were trypsinized, washed in PBS and resuspended in STE buffer (250mM sucrose, 5mM Tris and 2mM EGTA, pH 7.4) with 1% BSA. Cells were homogenized using 10 strokes with a tight fit dounce homogenizer. Cells were then centrifuged at 8500xg for 10 minutes at 4°C and the pellet was resuspended in STE with 1% BSA. Centrifugation at 700xg for 10 minutes at 4°C was done to get rid of nuclei. The nuclei pellet was discarded, and the supernatant was transferred to a new tube and centrifuged at 8500xg for 15 minutes at 4°C to pellet the crude mitochondrial fraction.

### RNA preparation and quantitative RT-PCR

mRNA was extracted from tissues or in vitro cultures with TRIzol reagent (Invitrogen) and purified using Direct-zol RNA miniprep (R2055, Zymo) following manufacture instruction with DNase treatment. Cytosolic RNA was extracted from in vitro murine brown adipocyte cells using the Cytoplasmic and Nuclear RNA purification Kit (21000, Norgen Biotek Corp.) following the manufacturer’s instructions with DNase treatment. Mitochondrial RNA was extracted from isolated mitochondria fraction. Briefly, the mitochondrial pellet was resuspended in STE buffer and treated with 7μL of RNase I (10U/μL) (EN0601, Thermo Scientific) for 5 minutes on ice to get rid of remaining non-mitochondrial RNA. The mitochondrial suspension was then centrifuged at 8500xg for 20 minutes at 4°C and the remaining pellet was resuspended in TRIzol reagent (15596018, Invitrogen) for downstream extraction of mitochondrial RNA using Direct-zol RNA miniprep (R2055, Zymo) with DNase treatment. cDNA was synthesized using Maxima First Strand cDNA synthesis kit (K1672, thermo scientific), and 10-20 ng cDNA was used for qPCR on a QuantStudio 5 real-time PCR system with either TaqMan Universal Master Mix II and validated PrimeTime primer probe sets (Integrated DNA Technologies), or Maxima SYBR Green/ROX qPCR Master Mix (2X) (K0221, Thermo Scientific) and designed primers. The ΔΔCt method was used to comparatively assess gene expression. Primer sequences listed in Table S1.

### RNA-sequencing

Murine brown adipocytes were treated with/without 4CBA (4mM) for 24 hr and n=3 independent biological replicates were pooled to generate total RNA samples. Sequencing libraries were constructed from mRNA using KAPA mRNA HyperPrep Kit (KK8580) and NEXFlex barcoded adaptors (Bioo Scientific). High-throughput sequencing was performed using a HiSeq 2500 instrument (Illumina) at the UC Berkely QB3 Core. Raw reads were mapped using STAR^60^ against mouse (mm10) genome. DESeq2^61^ was used to determine differential gene expression. Enriched transcription factor binding sites in gene promoters were identified using HOMER^62^. RNAseq data will be available for public at the Gene Expression Omnibus (GEO) under accession GSE165940.

### siRNA-mediated knockdown of target genes

Scramble or predesigned siRNA targeted to mouse ATF4, ATF5, CHOP or HRI (MISSION siRNA, MilliporeSigma) was diluted in Lipofectamine RNAiMAX reagent and applied on day 3 of differentiating adipocytes overnight (20 pmol of siRNA per one well of 12 well plate), and then cells were switched to a maintenance medium containing 10%FBS, P/S, insulin, T3 and rosiglitazone. 4CBA treatment occurred on day 5 to day 6 of differentiation.

### Plasma FGF21

Plasma collected using EDTA was used for FGF21 determination by Mouse/Rat FGF-21 Quantikine ELISA Kit (R&D System) per manufacture’s instruction.

### In Vivo Respirometry

Between 6 to 8-week-old male animals were fed a defined diet (Research Diets D12450J) under ambient temperature (23 °C) for two weeks. Food intake and body weight were monitored three times a week. Whole-body energy expenditure (VO2, VCO2) was recorded at indicated environmental temperatures using a Comprehensive Lab Animal Monitoring System (CLAMS, Columbus Instruments). Data were imported and analyzed using the web-based analysis tool for indirect calorimetry experiments (CalR)^63^.

### In Vitro Respirometry

Oxygen consumption rate (OCR) and extracellular acidification rate (ECAR) were measured in differentiated brown adipocytes using the Seahorse XFe24 Extracellular Flux Analyzer (Agilent). Cells were treated with 4CBA (4 mM) for 24 hours, and preincubated in the XF assay medium supplemented with 10 mM glucose, 2 mM sodium pyruvate, and 2 mM GlutaMAX at CO_2_ free incubator for an hour. Cells were subjected to the mitochondrial stress test by sequentially adding oligomycin (1μM), FCCP (1 μM), and antimycin/rotenone mix (1 μM /1 μM).

### Immunoblot

BAT mitochondria were isolated as previous described^2^ and protein lysates were extracted using RIPA lysis buffer. 15 ug of mitochondrial proteins or 25 ug of total cellular proteins were separated by gel electrophoresis on 4-20% gradient TGX gels and transferred onto PVDF or nitrocellulose membranes using Trans-Blot Turbo Transfer system (BioRad). Mitochondrial electron transport chain proteins were detected using Total OXPHOS Rodent WB Antibody Cocktail (ab110413). Proteins were detected with the indicated antibodies, and blots were images using Odyssey Imaging System. Image Studio Lite was used for protein quantification.

### UCP1 current measurement

Mitoplasts isolation and UCP1 current measurement were done as previous described^64^. Briefly, mitoplasts were isolated from BAT of C57/BL mice fed with a defined diet for 10-15 days. Patch pipettes were filled with 130 mM tetramethylammonium hydroxide (TMA), 1 mM EGTA, 2 mM magnesium chloride, 150 mM HEPES, (pH adjusted to 7.0 with D-gluconic acid, tonicity adjusted to ∼360 mmol/kg with sucrose). Whole-mitoplast UCP1 current was recorded in the bath solution containing 150 mM HEPES and 1 mM EGTA (pH adjusted to 7 with Trizma base, tonicity adjusted to ∼300 mmol/kg with sucrose). All experiments were performed under continuous perfusion of the bath solution. Data acquisition and analysis were performed using PClamp 10 (Molecular Devices) and Origin 7.5 (OriginLab). All electrophysiological data presented were acquired at 10 kHz and filtered at 1 kHz. Amplitudes of H+ currents were measured 25 ms after application of the −160 mV voltage step.

### Statistical analysis

Data were analyzed using Prism and are expressed as mean□±□SEM. Statistical significance was determined by unpaired two-tailed student t test or one-way ANOVA for the comparison of two conditions. Significance presented at P*<0.05, P**<0.01, and P***<0.001 compared to controls or otherwise indicated.

## Acknowledgements

This work was supported by NIH grants R01DK089202, R01DK118727 and R01DK126830 to A.S. G.A.T was supported by American Diabetes Association postdoctoral fellowship #1-19-PDF-058. We thank the UC Berkeley imaging core facility members Denise Schichnes, and Dr. Steve Ruzin for their support. We thank Dr. David L. Gasser for providing PDSS2-LoxP mice and Dr. Christopher M Adams for providing ATF4-LoxP mice.

## Author Contribution

Conceptualization, A.S. and C.C.; Methodology, C.C., P.H.Z., and G.A.T; Supervision, A.S. K.S. B.W. and Y.K.; Investigation and data analysis, C.C., A.G., G.A.T., and A.M.B.; Writing, A.S. and C.C.; Funding Acquisition: A.S. All authors read, edited and approved the manuscript.

## Declaration of Interests

The authors declare no competing interests.

Correspondence and requests for materials should be addressed to Andreas Stahl

## Supplementary Information is available for this paper

Table S1 rt-PCR primer sequence

Figure S1 Coenzyme Q_10_ supplementation in CoQ-deficient murine brown adipocytes Related to Figure1. (A) Mitochondrial complexes isolated from brown adipocytes were measured by western blot, n=3. (B) UCP1 gene expression of brown adipocytes treated with 4CBA with/without the supplementation of CoQ_10_ (2μM), n=6. (C) Upper panel: Representative UCP1-dependent H^+^ current recorded from whole brown fat mitoplasts before (black) or after CoQ10 treatment (20 μM). 1mM GDP (blue) was used as inhibitor of UCP1 activity. Lower panel: UCP1-dependent H^+^ current densities in the IMM of brown fat after CoQ10 treatment compared to as UCP1-dependent H^+^ current before treatment, n=5. Mean ± SEM.

Figure S2 Stress responses by CoQ deficiency

Related to Figure 2. (A-B) Gene Ontology (GO) analysis of RNAseq data from 48-h 4CBA treated brown adipocytes. Representative GO categories repressed or induced. (C) Mitochondrial morphology in brown adipocytes with treatment was visualized by confocal LSM880 FCS, scale bar 5 μm. (D) Phosphorylation of Drp1 in brown adipocytes was measured and quantified. (E-G) rtPCR analysis of genes in murine brown adipocytes treated with vehicle (CTL) or 4CBA for 7, 24, 48 and 72h, n=3. Data represents mean ± SEM

Figure S3 Targeting ISR-related components

Related to Figure 3. (A) *ATF4, CHOP*, and *ATF5* expression in brown adipocytes with 4CBA treatment after indicated siRNA knockdown of target genes (x-axis). (B) *UCP1* expression in 4CBA-treated brown adipocyte with siRNA-mediated knockdown of *CHOP* or *ATF5*. (C) *UCP1* and *HRI* gene expression in brown adipocytes transfected with scramble or HRI siRNA under vehicle control or 4CBA treatment for 24 hours. Data are mean ± SEM. Representative for three repeated experiments.

Figure S4 Metabolic parameters of PDSS2 knockouts

Related to Figure 4. (A) Oxygen consumption of PDSS2^FL^ and PDSS2^BKO^ animals at 4°C measured by CLAMS (n=4/group). (B) food consumption accumulated in two weeks. Data are mean ± SEM.

Figure S5 Quantitative PCR analysis of genes in tissues of PDSS2 knockouts

Related to Figure 5. (A) ISR-related genes in BAT of PDSS2 knockout PDSS2^BKO^ animals fed with HFD for 3 months, n=4/group. (B) *FGF21* gene expression in tissues of PDSS2^BKO^, n=4. (C) FGF21 protein levels in Liver or iWAT of PDSS2^BKO^, n=4. (D) stress-related gene expression in livers of liver-specific PDSS2 knockouts, n=4. (E) genes of UCP1-independent thermogenesis in BAT of BAT-specific PDSS2, ATF4 and PDSS2, or FGF21 and PDSS2 knockouts, n=6-8/group. (F) genes expression in iWAT of BAT-specific knockouts PDSS2^BKO^, DKO^ATF4^ and DKO^FGF21^, n=6-8/group. *,**,***p<0.05, 0.01, 0.001 compared to corresponding flox controls.

## Notes

### Competing Interest Statement

The authors have declared no competing interest.

