## Supplementary figures and images for "Coenzyme Q regulates UCP1 expression and thermogenesis through the integrated stress responses"

### Supplemental Figure 1

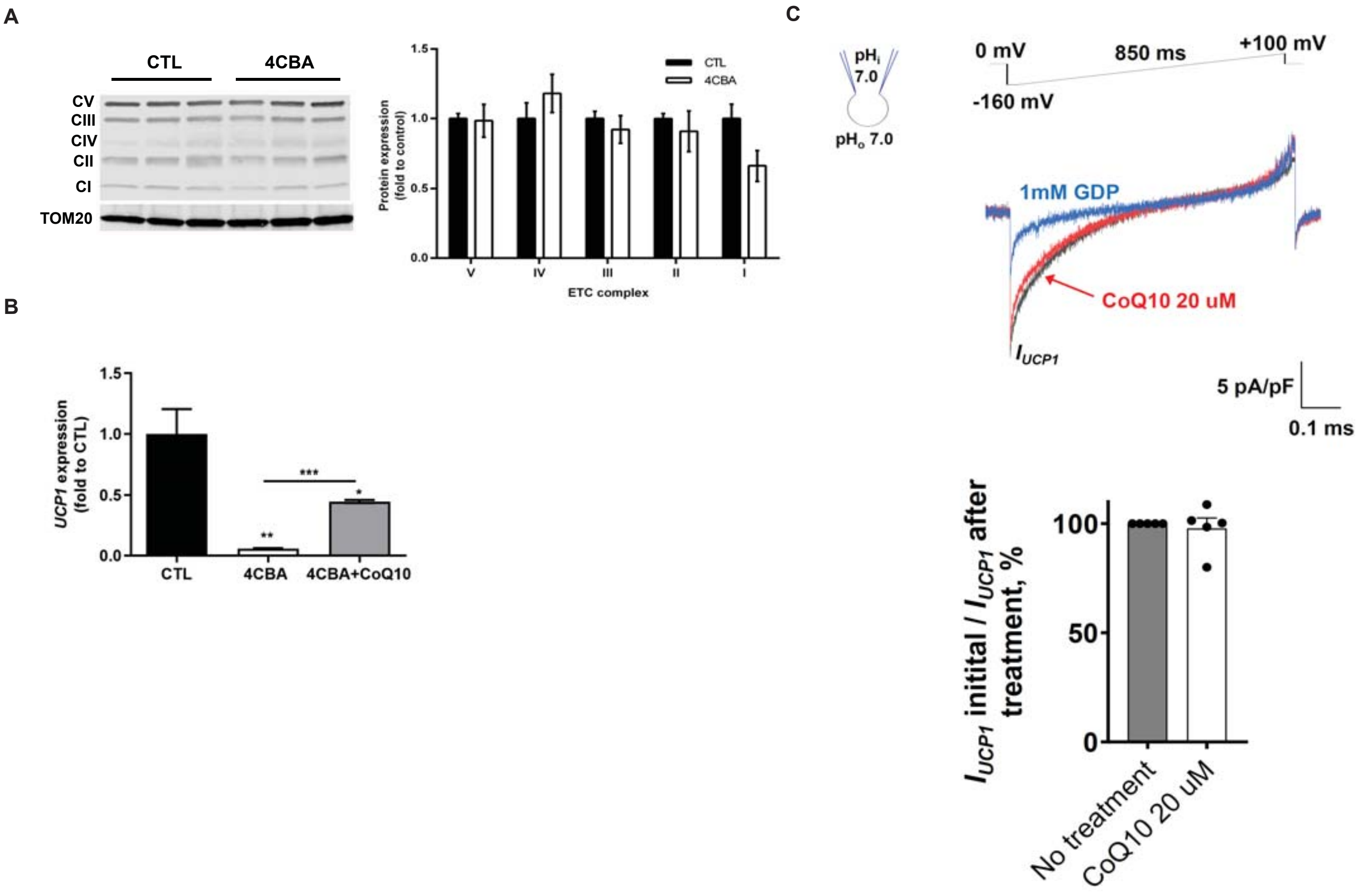

### Supplemental Figure 2

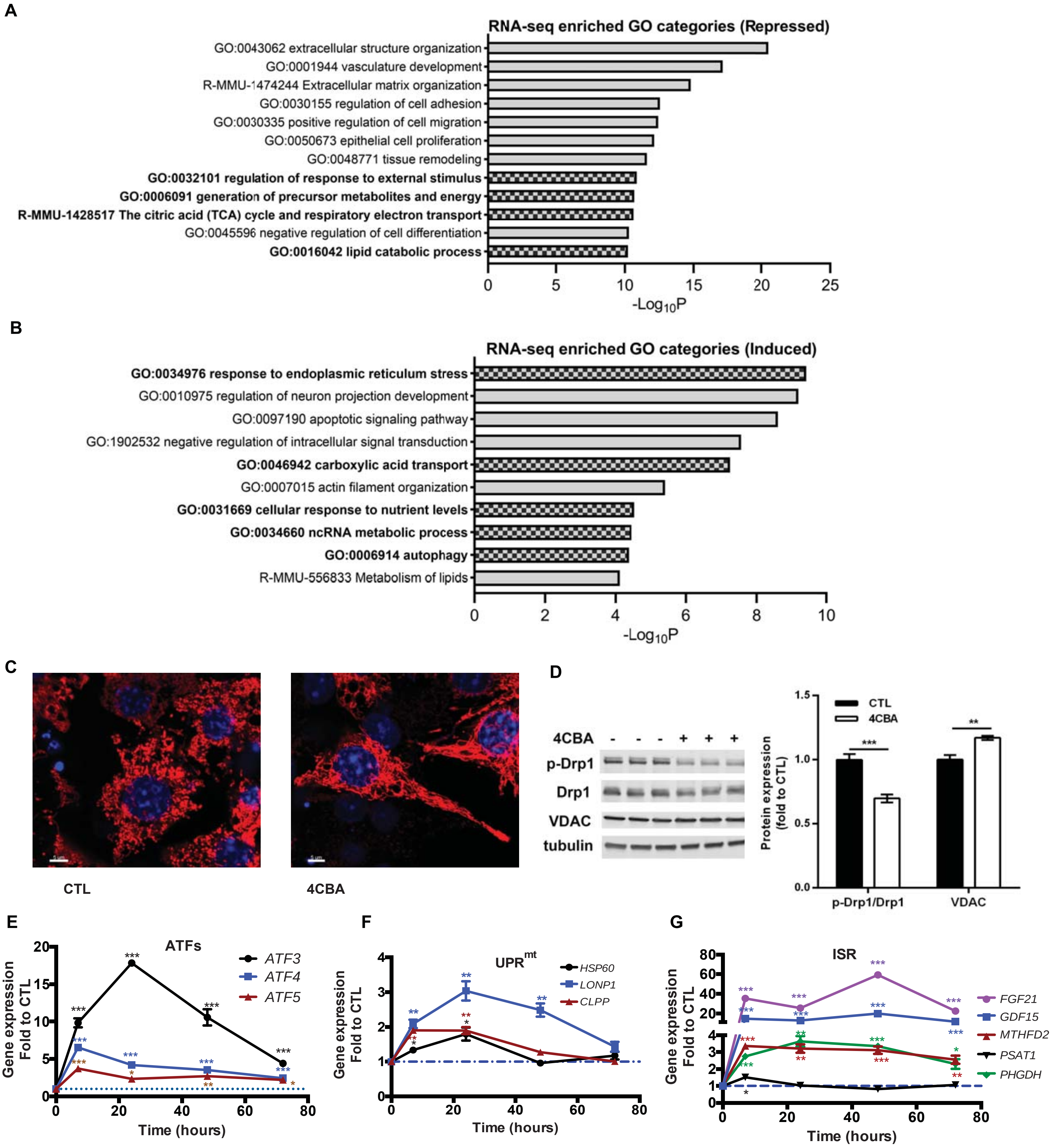

### Supplemental Figure 3

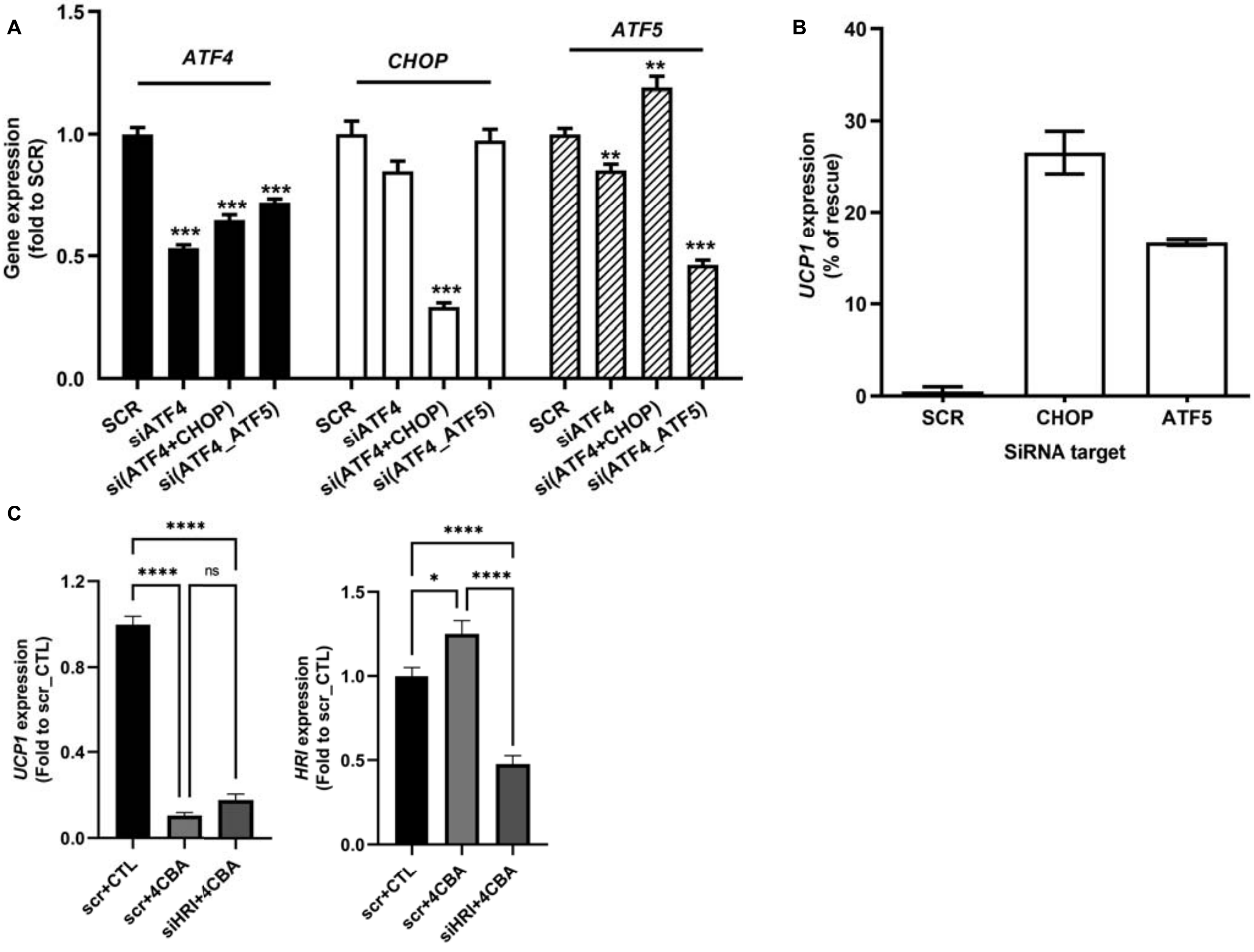

### Supplemental Figure 4

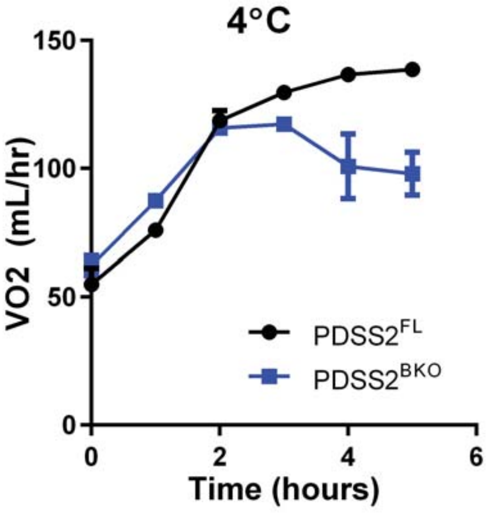

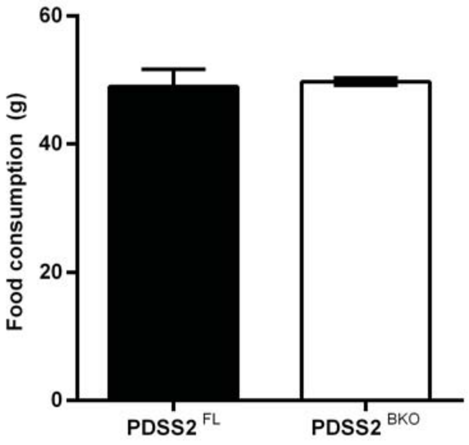

### Supplemental Figure 5

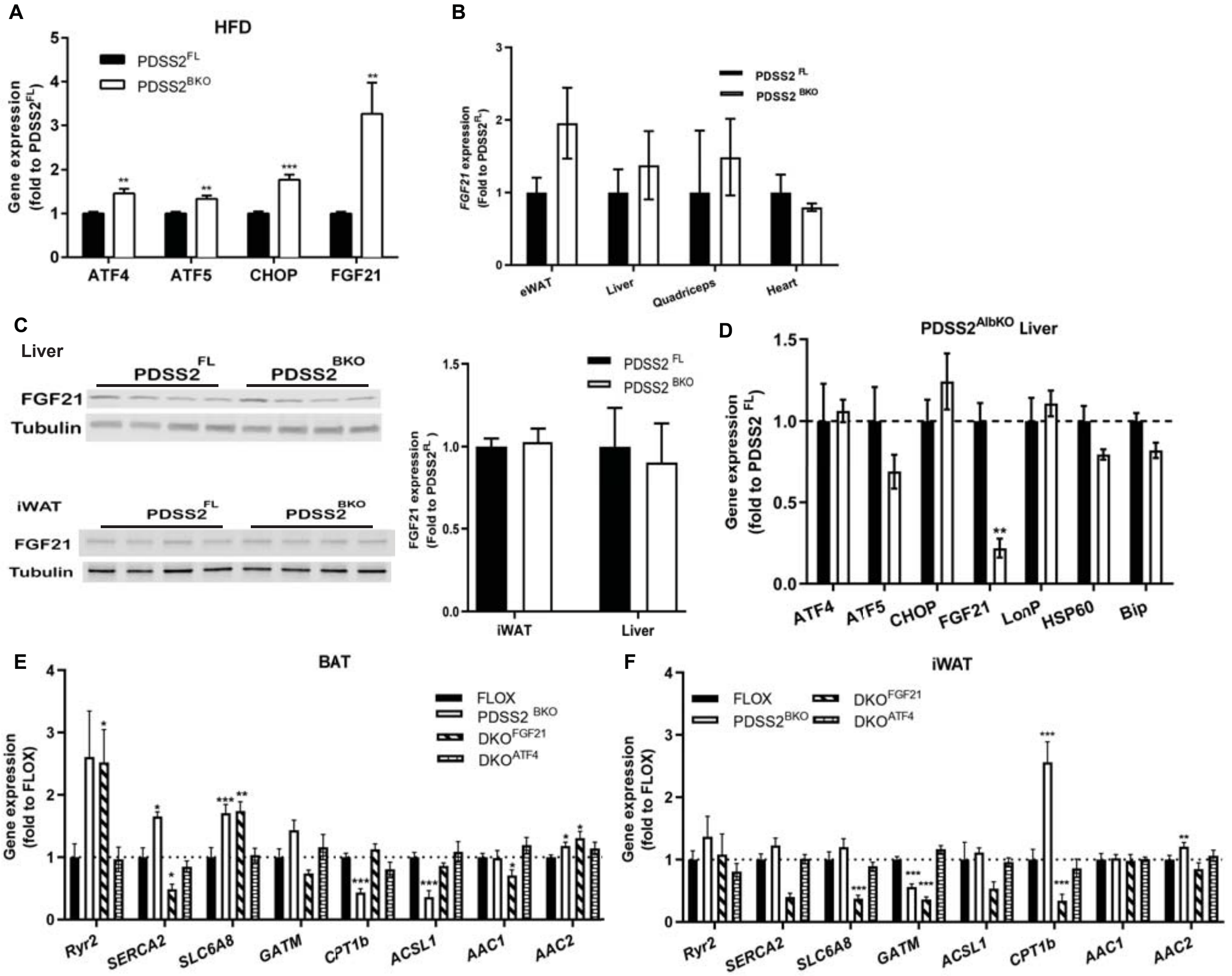
