## Supplemental Table 1 for "Coenzyme Q regulates UCP1 expression and thermogenesis through the integrated stress responses"

Table S1 related to Key Resources Table. rtPCR primer sequences.

| **Gene** | **Exons** | **Forward** | **Reverse** |
| --- | --- | --- | --- |
| UCP1 | 2-3 | CACACCTCCAGTCATTAAGCC | CAAATCAGCTTTGCCTCACTC |
| Mthfd2 | 2-3 | GTTCTCATCGTTGTTCAGCTTC | CGCCAGTCACTCCTATGTTC |
| Gdf15 | 1-2 | AGACCCTGACTCAGCGA | ACTCGAACTCAGAACCAAGTC |
| Psat1 | 2-3 | CCGAGTCCTCTGTAGTCTAGT | AGCTGTGCGGAATGAGATG |
| Phgdh | 8-9 | TTCTGCCAGACCAATCCAAG | AGTTTGTGGACATGGTGAAGG |
| ATF3 | 2-3 | TGTCTTCTCCTTTTTCTTGTTTCG | AGATGTCAGTCACCAAGTCTG |
| ATF4 | 1-2 | AGGTATCTTTGTCCGTTACAGC | CGTATTAGAGGCAGCAGTGC |
| ATF5 | 1-3 | AGCGTGGAAGATTGTTCAGC | GCTCACACCGTCTCTTCAG |
| CHOP | 1-3 | GACTCAGCTGCCATGACTG | GCGACAGAGCCAGAATAACAG |
| FGF21 | 1-1 | GGGATGGGTCAGGTTCAGA | CAGCCTTAGTGTCTTCTCAGC |
| PGC1a | 2-3 | CTCCATCTGTCAGTGCATCA | CCAACCAGTACAACAATGAGC |
| PRDM16 | 12-13 | CACTTGAACGGCTTCTCTTTG | CACAAGACATCTGAGGACACA |
| Ebf2 | 14-15 | GGAGGTGCTGTAATTAGATTGCT | GTCAGCATTTCAGAGTCCACA |
| Pparg | 4-5 | TGCAGGTTCTACTTTGATCGC | CTGCTCCACACTATGAAGACAT |
| Lonp1 | 10-11 | GGTTCTCTGTCTTGGTCTTCTTC | CTTCCGTTTCAGTGTTGGTG |
| Ppia | 4-5 | TTCACCTTCCCAAAGACCAC | CAAACACAAACGGTTCCCAG |
| PDSS2 | 6-8 | CACCTTTGCCAGCTTTGATTG | GTCTTACATCAGGAGTTTCTTGGA |
| HSP60 | 9-10 | TTTCTCATTCACTTCAACATCACTT | GCAGCTAGACATCACAACTAGTG |
| ERDj4 | 1-2 | AAGACGAAAACTGACTGTGGA | GAGGCTACTCGGCGTTC |
| Sdhb | 5-6 | TGTCTCCGTTCCACCAGTA | CAGTATCTGCAGTCCATCGAG |
| Tomm20 | 4-5 | ATTCCACATCATCTTCAGCCA | ACCAGTGTTCCAGATGCTTC |
| Clpp | 5-6 | TGTCCAAGATGCCAAACTCT | CAACATCTACGCCAAACACAC |
| PPIA |  | CAAACACAAACGGTTCCCAG | TTCACCTTCCAAAGACCAC |
| HK2 |  | ATTGTCCAGTGCATCGCGGA | AGGTCAAACTCCTCTCGCCG |
| SDHA |  | AGAAAGGCCAAATGCAGCTC | GTGAGAACAAGAAGGCATCAGC |
| ND1 |  | CTAGCAGAAACAAACCGGGC | CCGGCTGCGTATTCTACGTT |
| ND4 |  | CAACCAAACTGAACGCCTAAAC | GAGGGCAATTAGCAGTGGAATA |
| ND5 |  | CGGAGCCCTAACCACATTATT | GCCTAGTTGGCTTGATGTAGAG |
